# Effects of early geometric confinement on the transcriptomic profile of human cerebral organoids

**DOI:** 10.1101/2021.02.18.431674

**Authors:** Dilara Sen, Alexis Voulgaropoulos, Albert J. Keung

## Abstract

**Background:** Biophysical factors such as shape and mechanical forces are known to play crucial roles in stem cell differentiation, embryogenesis and neurodevelopment. However, the complexity and experimental challenges capturing such early stages of development, and ethical concerns associated with human embryo and fetal research, limit our understanding of how these factors affect human brain organogenesis. Human cerebral organoids (hCO) are attractive models due to their ability to model important brain regions and transcriptomics of early *in vivo* brain development. Furthermore, they provide three-dimensional environments that better mimic the in vivo environment. To date, they have been used to understand the effects of genetics and soluble factors on neurodevelopment. Establishing links between spatial factors and hCO development will require the development of new approaches.

**Results:** Here, we investigated the effects of early geometric confinements on transcriptomic changes during hCO differentiation. Using a custom and tunable agarose microwell platform we generated embryoid bodies (EB) of diverse shapes and then further differentiated those EBs to whole brain hCOs. Our results showed that the microwells did not have negative gross impacts on the ability of the hCOs to differentiate generally towards neural fates, and there were clear shape dependent effects on neural lineage specification. In particular, we observed that non-spherical shapes showed signs of altered neurodevelopmental kinetics and favored the development of medial ganglionic eminence-associated brain regions and cell types over cortical regions.

**Conclusions:** The findings presented here suggest a role for spatial factors in brain region specification during hCO development. Understanding these spatial patterning factors will not only improve understanding of *in vivo* development and differentiation, but also provide important handles with which to advance and improve control over human model systems for *in vitro* applications.

## BACKGROUND

Much of our understanding of brain development is and continues to be derived from foundational studies in animal models [1]. While these studies shed light into the general features of mammalian brain development, animal models do not capture the full complexity nor genetics of human neurodevelopment [2]. Recent progress with human cerebral organoid (hCO) systems show that these *in vitro* systems recapitulate many cell types and key features of the early *in vivo* brain making them attractive models to study human neurodevelopment [3–11]. As a model, they are continually being improved to better mimic *in vivo* architectures and brain regions and to enhance reproducibility, amongst other features. The development of any new understanding of factors regulating their development will therefore be important in advancing these models.

Human fetal development is controlled by cascades of differential gene expression patterns regulated by not only morphogens, but also spatial factors. Together these allow the formation of embryonic body axes and guide the generation and directionality of important structures such as the neural tube and primitive streak [12]. Furthermore, previous studies show that mechanical cues alone can bias stem cell behaviors and drive tissue patterning and multicellular organization [13–18]. Establishing links between mechanical cues and development is therefore an important step forward with regard to *in vitro* recapitulation of *in vivo* events, understanding how shape affects developmental changes, and improving organoid models [19].

Given the significant influence of spatial cues in directing human fetal development and hPSC differentiation, we hypothesized that shape may also influence hCO development and brain region specification and that these changes would be reflected in differential gene expression patterns. To test this, we engineered agarose microwells with distinct shapes but equivalent volumes and seeded embryoid bodies (EB) within them. The EBs conformed to these shapes in the first few days of culture, and we proceeded to expose them to a whole brain hCO protocol that is highly permissive of natural patterning due to minimal addition of exogenous growth factors [20]. We analyzed the transcriptomic changes in hCOs at different differentiation stages in both conventional 96-well plates and the agarose microwells by RNAseq to investigate the potential impacts of microwell culture and initial EB shape on neuronal differentiation or regional specifications. Intriguingly, shapes that conferred additional mechanical strain elevated general neuronal differentiation as well as forebrain and midbrain gene expression patterns. This effect was not immediately apparent in the first week but became clear as the hCOs matured for 3 weeks in culture. This work demonstrates that the shape, while holding total cell number constant, regulates neural microtissue differentiation and could be an important regulatory handle for hCO engineering and downstream mechanistic mechanobiology studies on human neurodevelopment.

## RESULTS

### Development and optimization of shape controlled hCOs in agarose microwells

The first step in the majority of current hCO protocols is the generation of embryoid bodies (EBs) and neuroectoderm from human pluripotent stem cells (hPSCs) [3, 21, 22]. Previous studies show that EBs are capable of developmental specification similar to that of the pre-gastrulation embryo and that the microenvironment and rigorous control of EB size strongly influence cell lineage-commitment [23–29]. In most hCO protocols, EB generation is carried out in low-attachment U- or V-bottom 96-well plates [20, 30] to form spherical aggregates of different sizes depending on the number of seeded cells.

While shape has previously been proposed as a potential factor regulating lineage commitment [13, 14, 28, 31], focus has been primarily on controlling EB size rather than shape especially in undirected differentiation protocols [24–29]. Here, we adapted a microengineering approach to fabricate distinctly shaped agarose microwells [32] (Figure 1A) (Additional file 1: Figure S1A, B). PDMS devices did not perform well with cells and organoids attaching to the device surfaces (Additional file 1: Figure S1C). We aimed to capture the first 5 days of EB generation after cell seeding in microwells, during which media conditions allow all germ layers to form spontaneously [6, 28]. Several design parameters such as volume and shape geometry were optimized for formation of a single EB in each microwell similar to those that generated in U-bottom 96-well plates (Figure 1B-D).

**Figure 1.**
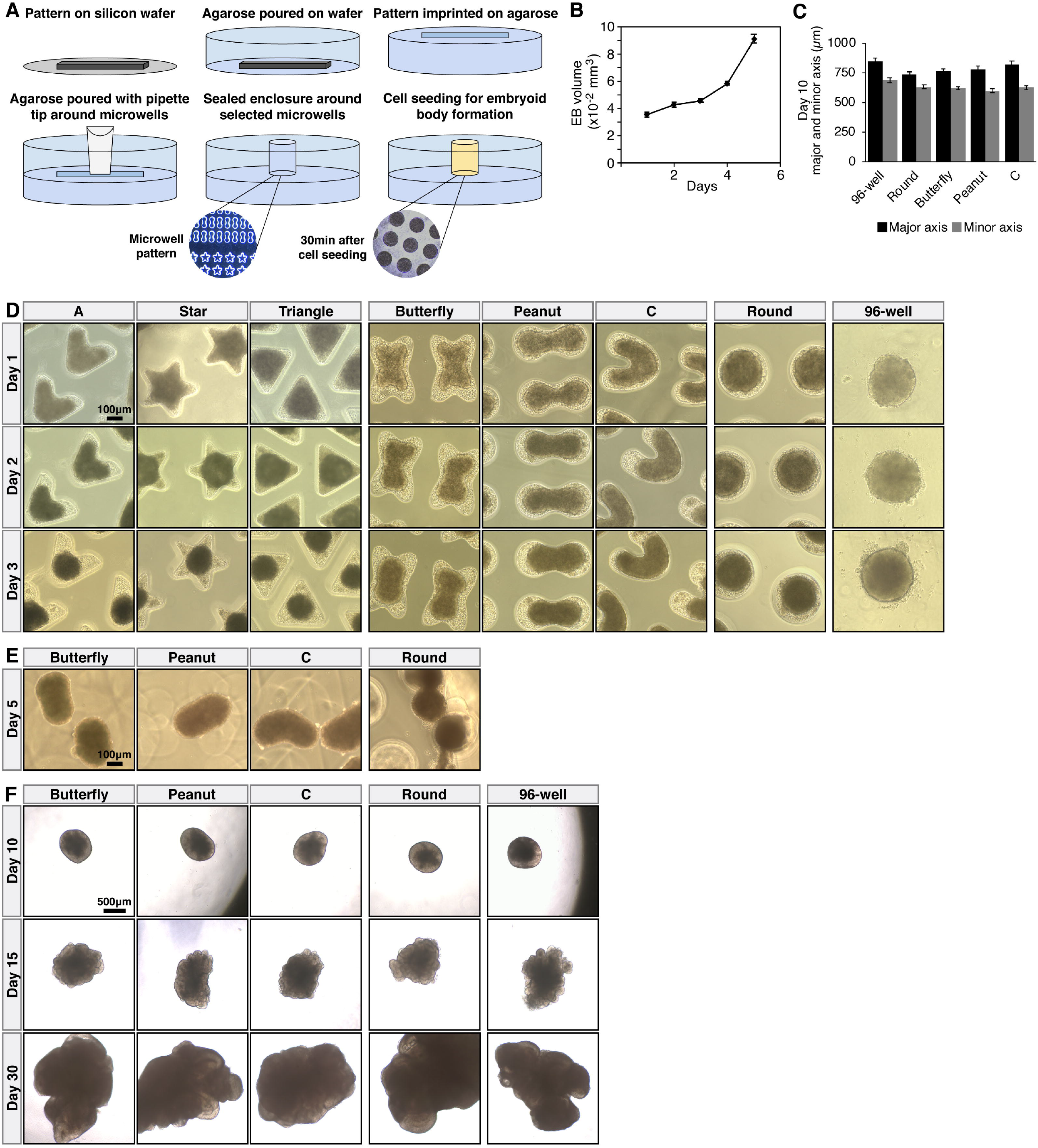
Engineering and optimization of early hCO shape-confinement in agarose microwells. A) Schematic description of the agarose microwell fabrication (see also Methods for details). B) Volume change of EBs seeded in a commercial 96-well plate. n=20 EBs per time point. Error bars = 95% confidence intervals. C) Size comparison of day 10 hCOs. n=50 hCOs per shape. Error bars = 95% confidence intervals. D) Development of EBs in distinctly shaped agarose microwells and 96-wells. E) Images showing the shape retention of EBs. F) Later developmental points of hCOs that were derived from EBs in microwells and 96-well plates.

To minimize any size-related effects and ensure the resulting EBs in microwells were comparable in size to the EBs generated in 96-wells, we first calculated the volume changes over 5 days for EBs generated in 96-well plates (Figure 1B). Then we adjusted the microwell design parameters and initial cell seeding density to achieve uniform EB sizes across all conditions (see also methods for details) (Figure 1C). Next, we tested the efficiency of forming EBs in several different geometries. While EBs successfully formed in almost all tested shapes after 24-hours, upon extended culture (3-4 days) we observed a ‘rounding effect’ in wells with high curvature (A, star and triangle), whereas the EBs in shapes with lower curvature (butterfly, peanut, C, round) retained the shape of the microwell for 4 days and even after 24-hours after removal from the wells (Figure 1D-E).

After 5 days, EBs were transferred from the microwell devices to low-attachment 24-well plates containing media driving neuroectoderm specification and hCO differentiation [20] (Figure 1F). Comparisons of major/minor axis ratios showed no significant differences in shape by day 10 of differentiation and all hCOs had adopted a similar spherical shape (Figure 1F) (Additional file 1: Figure S1D). In addition, all tissues showed smooth edges with bright optically translucent peripheries consistent with healthy neuroectoderm formation. There was also evidence of optically clear neuroephitelial bud outgrowth after Matrigel embedding (Figure 1F). These results indicated that hCOs initially seeded in shaped microwells (referred to as ‘microwell hCOs’ from here on) showed similar gross morphologies to hCOs developed in 96-well plates (referred as ‘96-well hCOs’ from here on), and they successfully met the developmental milestones of the whole brain hCO protocol followed in this study [20].

### Microwell and 96-well hCOs show similar general developmental capacities

Gross morphological and size characteristics indicated that microwell culture were not likely having adverse effects on the ability of hCOs to form and differentiate towards neural lineages. To further confirm this, we characterized the global transcriptomic changes associated with time in culture in all samples using RNAseq analysis (Figure 2). Principal component analysis (PCA) of samples from three time points (day 10, 20 and 40) identified notable transcriptional remodeling associated with differentiation with the greatest changes observed from day 10 to 40, and day 20 appearing as an expected transitional state (Figure 2A-B) (Additional file1: Figure S2A-B). Thus, for the remainder of our analysis we focused on the developmental changes between day 10 and day 40.

**Figure 2.**
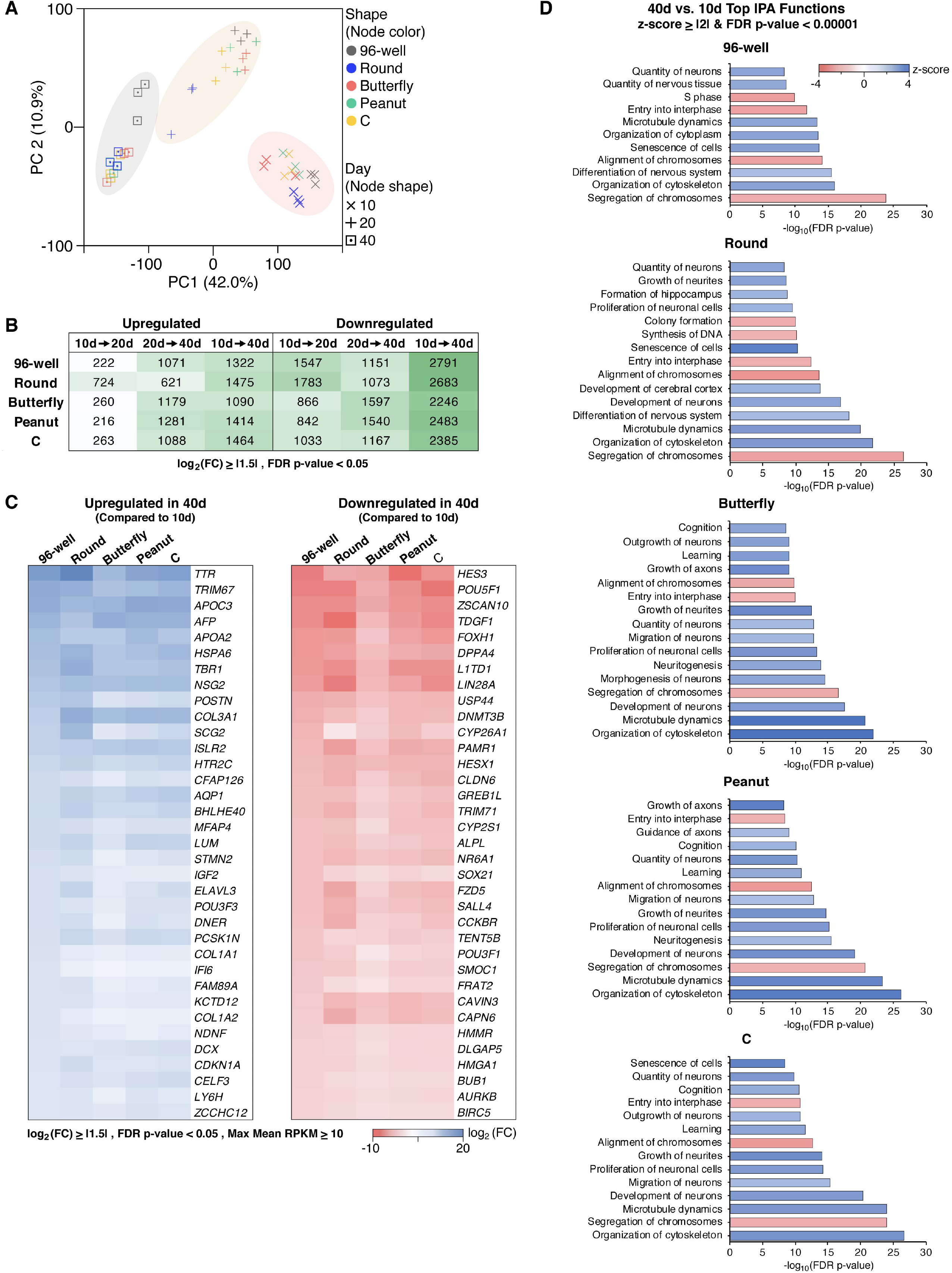
Microwell and 96-well hCOs show similar general developmental capacities. (A-D) n=3 biological replicates for each condition with 4-10 hCOs comprising each replicate. A) PCA of gene expression during hCO differentiation at days 10, 20 and 40. B) The number of differentially expressed genes during hCO differentiation with false discovery rate (FDR) p-value < 0.05 and log_2_ (fold change) (FC) ≥ |1.5|. C) The top 35 differentially expressed genes between days 10 and 40 with FDR p-value < 0.05, log_2_ (FC) ≥ |1.5| and maximum mean reads per kilobase of transcript per million mapped reads (max mean RPKM) ≥ 10. D) Top biological functions between days 40 and 10 with z-score ≥ |2| and FDR p-value < 0.00001 identified through IPA analysis. Differentially expressed genes with FDR p-value < 0.05, log_2_ (FC) ≥ |1.5| and max mean RPKM ≥ 10 were used in this analysis. 809 (96-well), 877 (Round), 621 (Butterfly), 807 (Peanut) and 788 (C) genes met this criteria.

We focused on the top differentially expressed genes in 96-well hCOs and determined the changes for the same list of genes in microwell hCOs (Figure 2C). This analysis revealed that in all samples as time in culture increased (day 40 versus 10) pluripotency associated genes (e.g., *HES3, POU5F1, ZSCAN10, TDGF1, DPPA3, L1TD1, LIN28A, USP44, DNMT3B, HESX1*) were downregulated, and neurodevelopment associated genes (e.g., *TTR, TRIM67, TBR1, NSG2, SCG2, ISLR2, HTR2C, AQP1, STMN2, ELAVL3*) were upregulated (Figure 2C).

This developmental trend was also supported by the Ingenuity Pathway Analysis (IPA) where all microwells and 96-well hCOs largely shared the same top IPA functions. While the primary functions predicted to be inhibited (z-score ≤ -2) were related to cell division (e.g., entry into interphase, alignment of chromosomes, synthesis of DNA), the functions predicted to be activated (z-score ≥ 2) were associated with neurodevelopment and cellular senescence (e.g., quantity of neurons, differentiation of nervous system, senescence) (Figure 2D) (Additional file 1: Figure S2C). Collectively, this analysis confirms that in addition to gross morphological similarities, transcriptional dynamics and overall differentiation programs during 96-well Hco development were conserved in microwell hCOs.

### Early microwell culture affects subsequent hCO neurodevelopment

While the microwell conditions were similarly supportive of overall hCO development as the 96-well conditions were, there were transcriptomic differences apparent even in the initial PCA analysis (Figure 2A). To identify potential changes associated with the microwell cultures in general, the transcriptomes of microwell hCOs were compared to 96-well hCOs at each time point. Resulting PCA analysis revealed clear transcriptomic differences between the microwell and 96-well environments (Figure 3A). To capture the overall differences, we analyzed the differentially expressed genes between all microwell (regardless of the specific geometry) and 96-well hCOs. Notably, although the differences were apparent at all time points, they were particularly clear at day 40 (Figure 3B).

**Figure 3.**
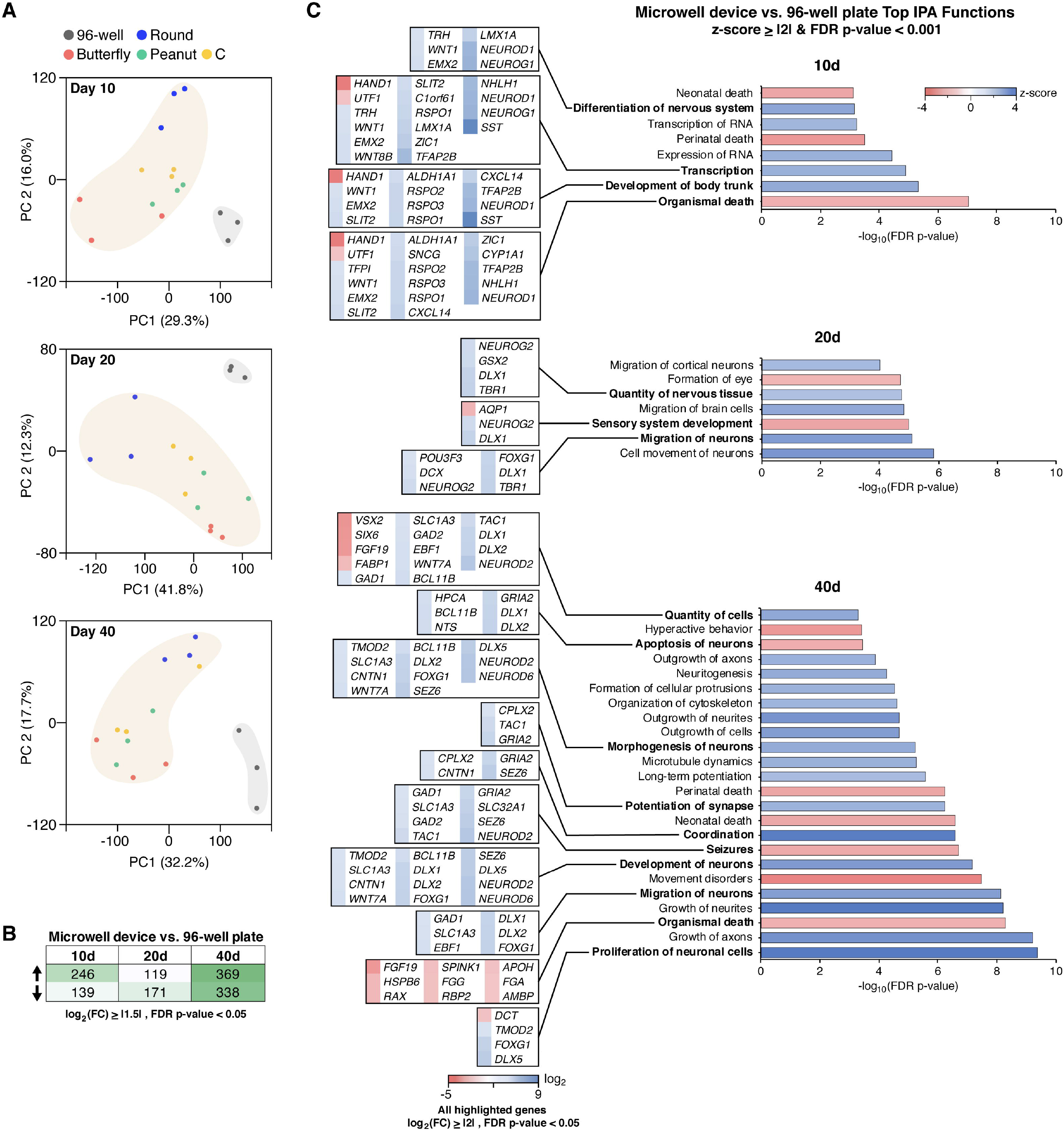
Early microwell culture affects subsequent hCO neurodevelopment. A) PCA of gene expression at individual time points. n=3 biological replicates for each condition with 4-8 hCOs in each replicate. (B, C) n=12 biological replicates for microwell device (all shapes) with 4-8 hCOs in each replicate, n=3 for 96-well plate with 4-8 hCOs in each. B) The number of differentially expressed (up arrow) upregulated and (down arrow) downregulated genes between microwell device and 96-well plate with FDR p-value <0.05 and log_2_ (FC) ≥ |1.5|. C) Top biological functions between microwell device and 96-well plate with z-score ≥ |2| and FDR p-value < 0.001 identified through IPA analysis. Differentially expressed genes with FDR p-value < 0.05 and log_2_ (FC) ≥ |2| associated with each function are highlighted. Differentially expressed genes with FDR p-value < 0.05, log_2_ (FC) ≥ |1.5| and max mean RPKM ≥ 5 were used in this analysis. 51 (day 10), 56 (day 20) and 166 (day 40) genes met this criteria.

Based on the differentially expressed genes, IPA results predicted an activation (z-score ≥ 2) of functions related to nervous system development (e.g., quantity of nervous tissue, morphogenesis of neurons, migration of neurons) (Figures 3C) (Additional file 1: Figure S3) in microwell hCOs at all three time points. Common inhibited (z-score ≤ 2) functions were associated to organismal death (Figure 3C) (Additional file 1: Figure S3). Functions related to sensory system development were also inhibited at days 20 and 40. Interestingly, while some functions associated with excitatory-inhibitory balance such as seizures were inhibited, others such as coordination were activated in day 40 microwell hCOs.

Notable transcriptomic differences associated with these biological functions were; the stark downregulation of mesoderm (*HAND1*, >5 fold) [33] and pluripotency (*UTF1*, >2 fold) [34] markers and the upregulation of dorsal medial ganglionic eminence (MGE) (*SST*, >8 folds) [7, 9, 10] and intermediate progenitor markers (*NEUROD1, NEUROG1, NHLH1, TFAP2B*, all >5 folds) [35, 36] at day 10, downregulation of retina associated genes (*VSX2, SIX6*, both >4 fold) [5] and upregulation of cortical (*NEUROD2, NEUROD6*, both >4 fold), forebrain (*FOXG1*, >3 fold) and ganglionic eminence (GE) (*DLX1, DLX2, DLX5, DLX6, SLC32A1*, all >3 fold) [7, 9, 10] markers at day 40 (Figure 3C). These intriguing differential gene expression patterns and IPA results indicate that the initial EB growth environment (microwell versus 96-well) may instruct a lineage bias in subsequent neural tissue development.

### Comparison of individual microwell geometries to 96-well hCOs reveal shape specific differences in gene expression

We then investigated how different microwell geometries contributed to the global transcriptomic differences observed in microwell and 96-well hCOs (Figures 4) (Additional file 1: Figure S4). Differential gene expression analyses between individual shapes and 96-well hCOs showed a varying trend of up and downregulated genes across the different geometries (Figures 4A) (Additional file 1: Figure S4A-C). This was particularly apparently at day 40 where round microwells showed an overall trend of downregulation, C microwells showed similar numbers of up and downregulated genes, and butterfly and peanut microwells were associated with upregulation of transcripts (Figures 4A) (Additional file 1: Figure S4C).

**Figure 4.**
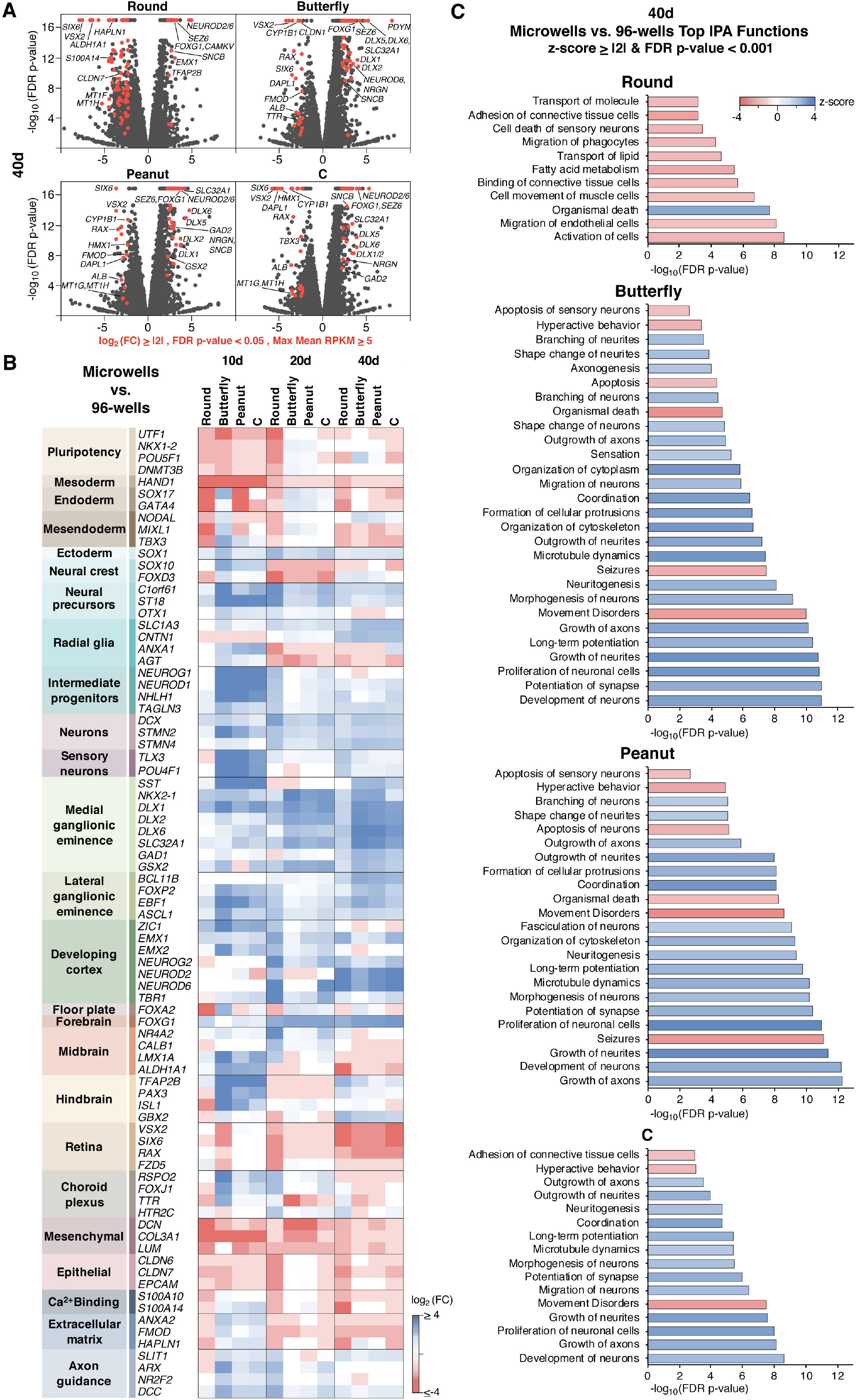
Comparison of individual microwell shapes to 96-well derived hCOs reveal shape specific differences in gene expression. (A-C) n=3 biological replicates for each condition with 4-10 hCOs in each replicate. A) Volcano plots of differentially expressed genes between individual microwell shapes and 96-wells at day 40. Red highlighted genes: FDR p-value < 0.05, log_2_ (FC) ≥ |2| and max mean RPKM ≥ 5. B) Differentially expressed marker genes between individual microwell shapes and 96-wells with FDR p-value < 0.05. C) Top biological functions between individual microwell shapes and 96-wells at day 40 with z-score ≥ |2| and FDR p-value < 0.001 identified through IPA analysis. Differentially expressed genes with FDR p-value < 0.05, log_2_ (FC) ≥ |1.5| and max mean RPKM ≥ 5 were used in this analysis. 174 (Round), 223 (Butterfly), 222 (Peanut) and 184 (C) genes met this criteria.

To gain a deeper understand of these differences, we constructed a panel of established cell type and brain region marker genes [4, 6, 7, 9, 10, 30, 36, 37] (Figure 4B). This analysis revealed that early EB growth in microwells instructed an early differentiation bias where genes associated with pluripotency and mesoderm were significantly downregulated in all microwell shapes at day 10. This was also accompanied by an upregulation in ectoderm marker *SOX1*. All microwell shapes showed downregulation of neural crest markers and upregulation of neural progenitor markers at day 20. Interestingly markers associated with intermediate progenitors and sensory neurons were only significantly upregulated in non-spherical shapes (butterfly, peanut and C). In addition, these non-spherical geometries also showed a drastic upregulation of MGE markers *SST* and *DLX1* at day 10 followed by a robust upregulation of all MGE and lateral ganglionic eminence (LGE) associated genes at day 40. While some cortical markers (*NEUROD2, NEUROD6*) showed significant upregulation at day 40 in all shapes, a notable trend for cortical development was not observed across different shapes and time points. Similarly, floor plate, midbrain and hindbrain associated genes also showed varying results, whereas forebrain marker *FOXG1* demonstrated significant upregulation in all microwell shapes at day 20 and 40. Furthermore, epithelial and mesenchymal fates and retinal development genes showed a notable downregulation in microwell hCOs (day 20 & 40 for epithelial and retina, all time points for mesenchymal markers).

These differences between microwells and 96-well and across different microwell geometries were also supported by IPA analyses (Figure 4C) (Additional file 1: Figure S4D-E). In agreement with the differential expression analysis, while round microwells showed an overall tendency for inhibition (z-score ≤ 2), the biological functions in butterfly, peanut and C were mostly associated with activation (z-score ≥ 2) at day 40 (Figure 4C). These results demonstrate that while there are common features affected in all geometries indicating they could be the result of microwell growth environment, there are also differences specific to distinct geometries implying shape-associated transcriptomic differences in hCOs.

### Microwell geometry modulates gene expression programs related to neural cell type and brain region specification

Finally, we shifted our focus to the transcriptomic changes across different microwell geometries to investigate whether shape was informative of brain region specification (Figure 5) (Additional file 1: Figure S5). To isolate the shape specific effects from differences arising from microwell devices in general, round microwells were used as the control group in this analysis as this shape is the closest to the naturally occurring spherical shape of an EB.

**Figure 5.**
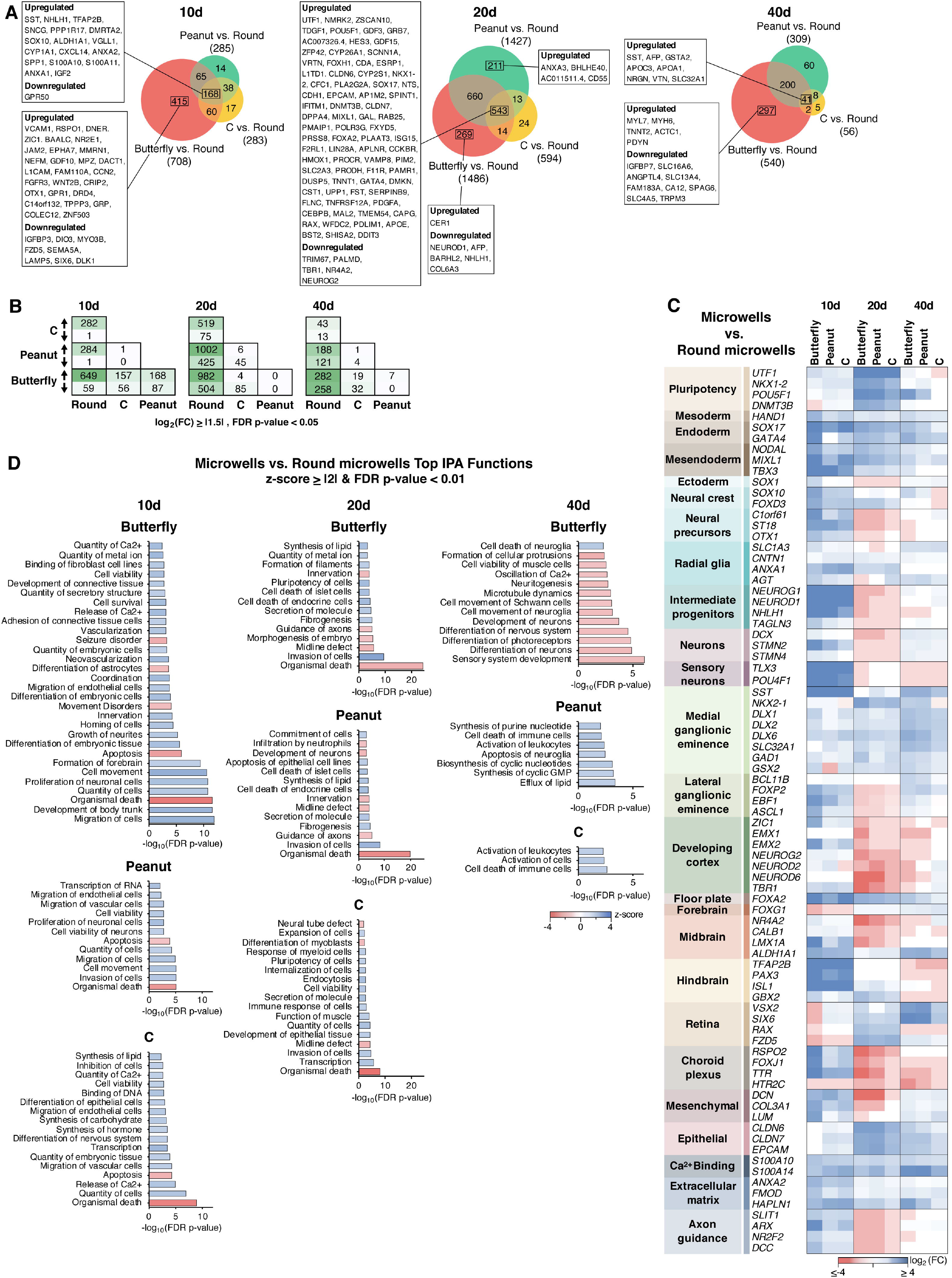
Microwell geometry modulates gene expression programs related to neural cell type and brain region specification. (A-D) n=3 biological replicates for each condition with 4-10 hCOs in each replicate. A) Venn diagrams showing the differentially expressed genes with FDR p-value < 0.05 and log_2_ (FC) ≥ |1.5|. Boxed genes are the unique and common transcripts in each comparison with FDR p-value < 0.05, log_2_ (FC) ≥ |2| and max mean RPKM ≥ 5. B) The number of differentially expressed genes across all microwell geometries with FDR p-value <0.05 and log_2_ (FC) ≥ |1.5|. C) Differentially expressed marker genes between individual microwell geometries and round microwells with FDR p-value < 0.05. D) Top biological functions between individual microwell geometries and round microwells with z-score ≥ |2| and FDR p-value < 0.01 identified through IPA analysis. Differentially expressed genes with FDR p-value < 0.05, log_2_ (FC) ≥ |1.5| and max mean RPKM ≥ 5 were used in this analysis. 149 (Butterfly), 28 (Peanut) and 35 (C) genes met this criteria in day 10 hCOs. 286 (Butterfly), 266 (Peanut) and 88 (C) genes met this criteria in day 20 hCOs. 106 (Butterfly), 69 (Peanut) and 10 (C) genes met this criteria in day 40 hCOs.

Differential expression analysis revealed that while C microwells had the lowest number of differentially expressed genes at all time points, butterfly microwells showed the highest (Figure 5A,B). Pairwise comparisons between all shapes demonstrated that peanut and C microwells did not show considerable transcriptomic differences from each other (Figure 5B). While earlier differences observed between butterfly and peanut microwells were diminished at later points of differentiation, all shapes showed marked deviation from the round microwells at all time points (Figure 5B).

The marker gene panel revealed several interesting differences between non-spherical shapes and round microwells. First, we noticed a robust upregulation at day 10 and a downregulation at day 20 for markers associated with neural differentiation (neural precursors, intermediate progenitors and neurons) which applied to all shapes (Figure 5C). To understand the basis of this pattern, we re-examined our previous comparison between distinct microwell shapes and 96-well hCOs (Figure 4B). This analysis revealed that while butterfly, peanut and C microwells showed upregulation in these markers at day 10, the same change was not observed in round microwells until day 20 (Figure 4B) which may imply that the neural differentiation timeline is lengthened in round microwells (or accelerated in non-spherical shapes). A similar effect was also observed for LGE and axon guidance markers.

There were also cell type and brain region specific differences between shapes (Figure 5C). Notably, we observed an overall downregulation of cerebral cortex, forebrain, midbrain, choroid plexus and mesenchymal markers in non-spherical shapes (Figure 5C). In contrast, MGE, floor plate, hindbrain, epithelial and extracellular matrix markers were upregulated in these shapes (Figure 5C). These results may indicate that while round microwells promote cortical fates, other shapes favor MGE-associated brain regions and cell types. Overall, the most drastic changes were observed in butterfly microwells, followed by peanut and C geometries.

Supporting these observations, IPA results also highlighted butterfly microwells as the shape that was associated with the most significant alterations in biological functions (Figures 5D) (Additional file 1: S5). In addition, IPA results also revealed that there was an overall inhibition (z-score ≤ 2) of cell death related functions (e.g., organismal death, apoptosis) and activation (z-score ≥ 2) of cell viability linked functions (e.g., quantity of cells, cell viability, cell viability of neurons) in non-spherical shapes (Figure 5D).

## DISCUSSION

During the past decade, various *in vitro* strategies have been developed to study human brain development [38]. Among these, recently developed hCO systems are particularly advantageous due to their ability to capture important brain regions and cell types [2, 3]. While the primary focus in the field has been on identifying appropriate soluble factor cocktails to drive hCO formation, the present work provides a pilot analysis dedicated to the evaluation of geometric confinement effects on hCO development.

Our analysis comparing the microwell growth environment to 96-wells (Figure 3 & 4) showed apparent differences between transcriptional profiles including the stark downregulation of pluripotency markers, mesoderm, retina associated genes and mesenchymal fates in all microwell hCOs. Although, the microwell parameters were optimized with rigorous calculations and experimentation of several different conditions, these overall changes could be due simply to the confined geometric environment in the microwells compared to the unconfined U-bottom 96-well plates. In addition, unlike the U-bottom 96-well plates, the microwells had flat bottoms due to the fabrication method. While concave microwells have been used by others [24], their effects on differentiation in comparison to flat-bottomed wells, and in the context of shape, have not been investigated. Future studies could revise the fabrication methods used in this study to fabricate concave microwells in non-circular shapes to eliminate the potential transcriptomic differences that could result.

All of the non-spherical shapes investigated in this study showed similar patterns of differential gene expression compared to round microwells (Figure 5). However, these changes were more pronounced in butterfly shaped microwells, followed closely by the peanut microwells. C shape microwells showed the closest transcriptional profile to the round microwells. Interestingly, this pattern correlates with the number of strain or stress points applied to EBs in each geometry. Future work could fine tune these shapes to achieve uniformity across all other parameters and specifically control for the number of stress-strain points. Comparing similar geometries with slightly different parameters such as size, curvature, and major over minor axis ratios could help elucidate the specific shape-related bases of these transcriptional differences.

Among the differences we observed across shapes, the stark upregulation of dorsal MGE marker *SST* at day 10 followed by other hallmark MGE markers such as *NKX2-1, DLX1*/*2, GAD1* and *SLC32A1* (also known as *VGAT*) were the most notable changes. Coupled with the downregulation of cortical markers such as *TBR1, NEUROD2/6* and *EMX1/2*, this pattern in non-spherical shapes resembled the previously described MGE [7] and ventral forebrain hCOs [9, 10]. Interestingly, while the whole brain hCO differentiation protocol followed in this study [6, 20] enables the generation of multiple different brain regions, dorsal cortical regions are more frequently identified compared to ventral fates [6, 38, 39]. To increase the consistency and yield of ventral regions, previous studies modified this protocol with specific drug treatments [9]. Although further validation studies with immunohistochemistry would be needed to confirm these results, the transcriptomic changes observed in the microwell growth environment and distinct shapes may imply that in addition to specific signaling factors, early geometric confinement or mechanical perturbations could be another method to regulate brain region specification.

## CONCLUSIONS

Here, we used agarose microwells to control EB shapes and generated whole brain hCOs from these EBs to investigate the effects of early spatial regulation on transcriptomic changes during brain region specification.

Our results showed that gross morphological and transcriptional changes related to hCO differentiation were similar between 96-well and microwell hCOs, indicating that the microwell growth environment did not significantly affect the overall neural differentiation or viability of the hCOs (Figure 1 & 2). However, investigating individual developmental time points and comparing across 96-well and microwell conditions more closely revealed notable differences in genes associated with pluripotency, mesoderm, MGE, retina and cortical development, and epithelial and mesenchymal fates (Figure 3), implying that the microwell growth environment may bias lineage commitment in neurodevelopment. Furthermore, we found that the direction and intensity of this transcriptional bias was associated with the starting EB shapes (Figure 4 & 5). The butterfly shape showed the most noticeable differences, followed closely by the peanut shape. Among other notable differences, round microwells exhibited a potential delay in the kinetics of neurodevelopment, and our results also demonstrated that non-spherical shapes may favor the development of MGE-associated brain regions and cell types over cortical regions (Figure 5).

The findings presented here suggest a role for mechanobiological factors in brain region specification and hCO development. Uncovering how fate decisions are tied to these early spatial changes could provide a better understanding of how embryonic morphogenetic processes are orchestrated. In addition, we believe that these results not only offer a new perspective to advance our understanding of human brain development, but also pave the way for new experimental directions to refine the engineering of *in vitro* model systems and organoids.

## METHODS

### Agarose microwell device fabrication

Shapes were designed using Autocad (Autodesk) and photomasks were printed using Fineline Imaging printing services. The silicon wafer with these shapes was fabricated using photolithography [40]. Briefly, molds were fabricated by coating SU-8 2150 photoresist with a Laurell spin coater at 500 rpm for 7 seconds then at 1000 rpm for 35 seconds. Following photolithography, agarose microwells were fabricated by replica-molding (Figure 1A)

(Additional file 1: Figure S1A-B) [32]. Briefly, using a 10 cm circular stencil, a 1 cm layer of 2% agarose in PBS was poured onto the silicon wafer. Following this, air bubbles were removed by degassing and agarose was cooled at room temperature for 30 minutes to allow for congealing. Then, the agarose disc was gently removed and placed into a sterile 6-well culture plate, with the imprinted patterns facing upward. A cut pipette tip was placed on the pattern, so that roughly 80 patterned microwells were enclosed in the tip. With the pipette tip in place, warm agarose was poured in the remainder of the well. After the second layer of agarose cooled and congealed, the pipette tip was removed, leaving a small enclosure with shaped microwells (Figure 1A) (Additional file 1: Figure S1B). Then, 2 mL of 1x PBS was added to the agarose device and it was sterilized under UV light overnight. Base surface area and height of microwells were 0.117 mm^2^ and 0.6 mm, respectively, to achieve a 0.07 mm^3^ total volume in the microwells.

### Cell culture and human cerebral organoid generation

Feeder-independent H9 human embryonic stem cells (hESCs) (WA09) were obtained from WiCell. Cells were maintained in tissue culture dishes (Fisher Scientific Corning Costar) coated with 0.5 mg/cm2 Vitronectin (VTN-N) (Thermo Fisher Scientific) in E8 medium (Thermo Fisher Scientific) and passaged using standard protocols. 96-well and microwell human cerebral organoids (hCOs) were generated using the same pool of hESCs and differentiated using the same protocol as described [6, 20]. Number of cells seeded per well was experimentally optimized for successful generation of EBs in microwells. For each microwell device, 60 μL of hESC suspension at 32,000 cells/well was pipetted into the microwell enclosure using a cut p200 tip. The device was gently placed into the incubator for 5 minutes to allow cells to settle into the microwells. After 5 minutes, 2.5 mL media was slowly added to the wells avoiding the disruption of the seeded cells in the microwells. Media was changed every 12 hours for the first 3 days with fresh hESC medium with basic fibroblast growth factor (bFGF) (Invitrogen) and ROCK inhibitor (Y-27632, LC laboratories) for all hCOs (microwell and 96-well). All cells and hCOs were maintained in a humid incubator at 37 °C with 5 % CO_2_. After 5 days all hCOs were transferred into low-attachment 24-wells plates (1 hCO/well) and the remainder of the previously described whole brain organoid protocol was followed [6, 20].

### Total RNA extraction and sequencing

For RNA extraction the following samples were collected from three independent culture plates: 10 pooled hCOs for day 10 samples, 6 pooled hCOs for day 20 samples, and 4 pooled hCOs for day 40 samples. hCOs were washed 3 times in cold PBS. Total RNA was extracted as previously described [41]. Briefly, Matrigel was dissolved by incubating the hCOs in chilled Cell Recovery Solution (Corning, cat. no. 354253) for 1h at 4 °C. The dissolved Matrigel was removed by rinsing 3 times in cold PBS. Total RNA was isolated using Direct-zol RNA MicroPrep Kit (Zymo Research) according to the manufacturer’ s protocol. RNA samples were collected in 2mL RNAse-free tubes and chilled on ice throughout the procedure. Bulk RNA was sequenced using DNB-seq PE100 platform (BGI).

### RNAseq analysis

Raw FASTQ formatted sequence reads were imported into CLC Genomics Workbench (version 20.0.2, QIAGEN Digital Insights). Adaptor sequences and bases with low quality were trimmed and reads were mapped to the reference genome (GRCh38.102) using the RNAseq analysis tool with the default parameters recommended for RNAseq analysis. Principal component analysis and differential expression analysis were performed using ‘PCA for RNAseq’ and ‘Differential Expression for RNA-seq’ toolsets. ‘Wing’ like patterns observed in volcano plots reflect the mathematical relationship between fold change and p-values. This was exposed in our data set due to the number of replicates that were used, and the genes that appeared in these regions were not included in the analyses. Heatmaps were generated using Euclidean distance with complete linkage. The top 10,000 features with highest coefficients of variance were displayed. Significance was selected as log_2_(fold change) ≥ |1.5| in expression between comparison groups with a threshold false discovery rate (FDR) adjusted p-value < 0.05. All sequencing data passed default quality filters for the CLC Genomic Workbench version 20.0.2 RNA-Seq pipeline analysis (sequencing statistics summary available upon request). FASTQ sequencing files and sample metadata will be made publicly available through the National Center for Biotechnology Information Sequence Read Archive.

Ingenuity Pathway Analysis (IPA, version 60467501, QIAGEN Digital Insights) was used to predict activated or inhibited biological functions. Differentially expressed genes with log_2_(fold change) ≥ |1.5|, FDR p-value < 0.05, maximum mean reads per kilobase of transcript per million mapped reads (max mean RPKM) ≥ 10 (Figure 2) and max mean RPKM ≥ 5 (Figures 3, 4, 5) were used in IPA. Statistical z-score (≥ 2 for activation and ≤-2 for inhibition) was used to identify the predicted activity status of significant (FDR p-value < 0.05) biological functions. Cancer related biological functions were excluded from the analyses.

## ADDITIONAL FILES

Additional file 1: Supplementary figures S1-5. Extended data related to main figures 1-5. (PDF 9.4 mb)

## ETHICS APPROVAL AND CONSENT TO PARTICIPATE

Not applicable.

## CONSENT FOR PUBLICATION

Not applicable.

## AVAILABILITY OF DATA AND MATERIALS

The datasets generated and/or analyzed during the current study are available from the corresponding author upon reasonable request.

The accession number for the bulk RNA seq reported in this paper will be made publicly available.

## AUTHORS’ CONTRIBUTIONS

DS, AV and AJK conceived the study. DS and AV planned and performed the wet lab experiments with guidance from AJK. DS performed the RNAseq analysis. DS and AJK wrote the paper.

## ACKNOWLEDGEMENTS

We thank Drs. Adriana San Miguel and Sahand Saberi-Bosari for their valuable training in microfabrication process and allowing us to use their equipment.

