## Supplemental figures for "Effects of early geometric confinement on the transcriptomic profile of human cerebral organoids"

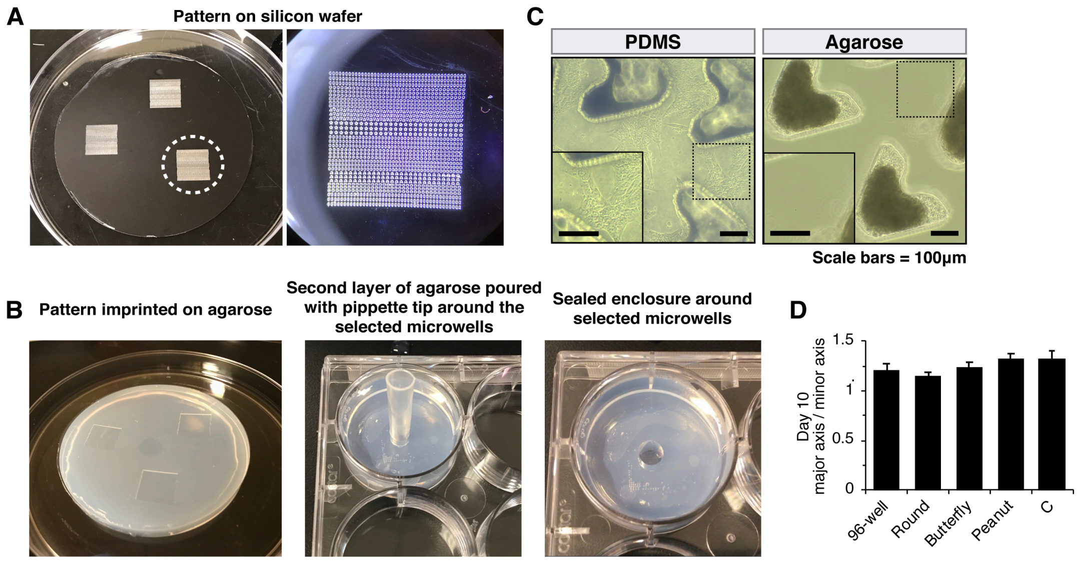


**Figure S1.** ***Related to Figure 1*****. Microwell device fabrication process and hCO size comparison at day 10.**

A) Images from silicon wafer mold (right) Image of the dotted outline area from (left).

B) Images from agarose microwell fabrication process.

C) Major/minor axis ratios of 10 day hCOs. n=50 hCOs per shape. Error bars = 95% confidence intervals.

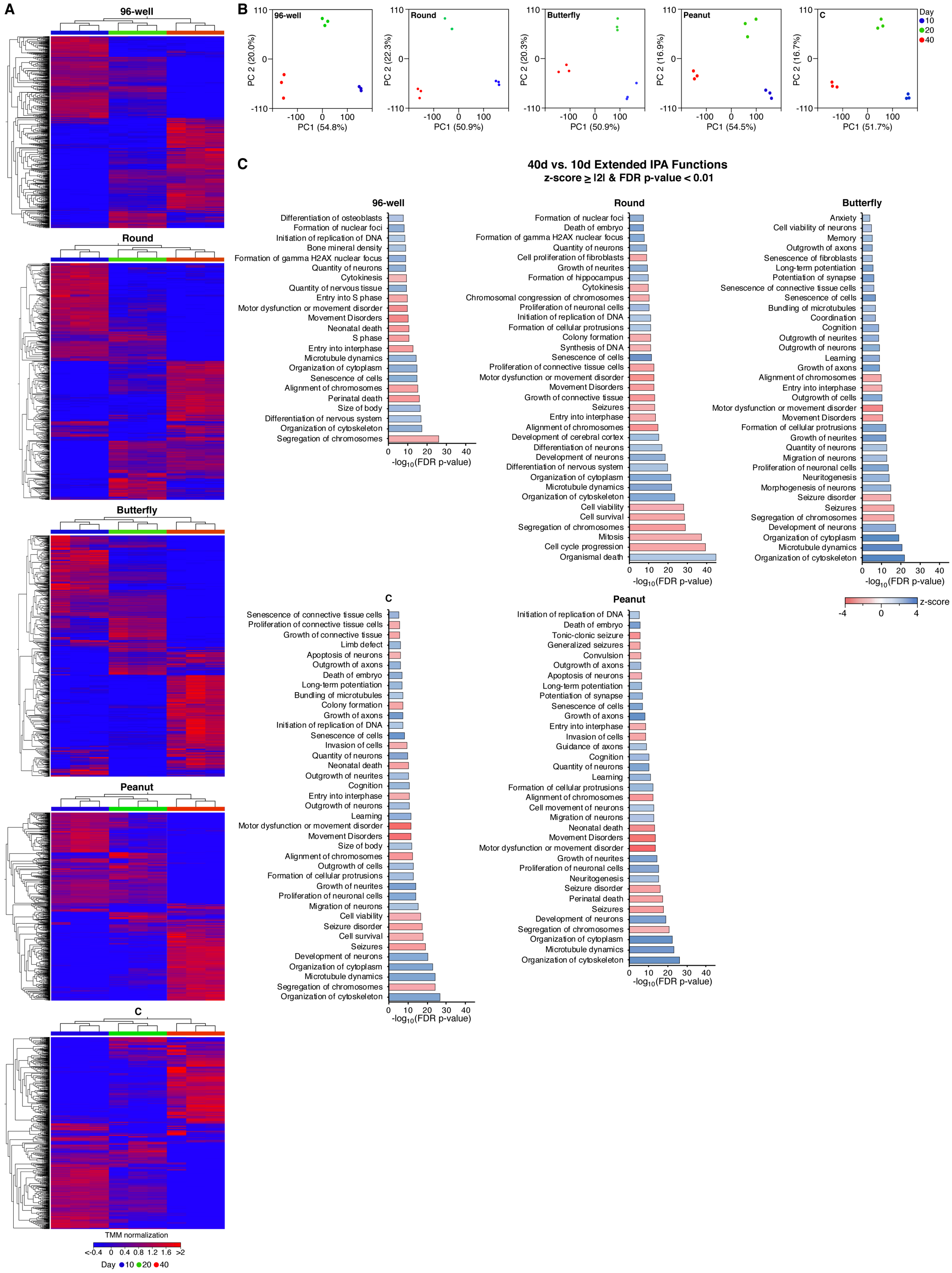
A
96-well
Round
Butterfly
Peanut
C
TMM normalization
<-0.4 0 0.4 0.8 1.2 1.6 >2
Day 10 20 40
B
96-well
Round
Butterfly
Peanut
C
PC1 (54.8%)
PC1 (50.9%)
PC1 (50.9%)
PC1 (54.5%)
PC1 (51.7%)
PC 2 (20.0%)
PC 2 (22.3%)
PC 2 (20.3%)
PC 2 (16.9%)
PC 2 (16.7%)
Day
10
20
40
C
40d vs. 10d Extended IPA Functions
z-score ≥ |2| & FDR p-value < 0.01
96-well
Differentiation of osteoblasts
Formation of nuclear foci
Initiation of replication of DNA
Bone mineral density
Formation of gamma H2AX nuclear focus
Quantity of neurons
Cytokinesis
Quantity of nervous tissue
Entry into S phase
Motor dysfunction or movement disorder
Movement Disorders
Neonatal death
S phase
Entry into interphase
Microtubule dynamics
Organization of cytoplasm
Senescence of cells
Alignment of chromosomes
Perinatal death
Size of body
Differentiation of nervous system
Organization of cytoskeleton
Segregation of chromosomes
-log10(FDR p-value)
Round
Formation of nuclear foci
Death of embryo
Formation of gamma H2AX nuclear focus
Quantity of neurons
Cell proliferation of fibroblasts
Growth of neurites
Formation of hippocampus
Cytokinesis
Chromosomal congression of chromosomes
Proliferation of neuronal cells
Initiation of replication of DNA
Formation of cellular protrusions
Colony formation
Synthesis of DNA
Senescence of cells
Proliferation of connective tissue cells
Motor dysfunction or movement disorder
Movement Disorders
Growth of connective tissue
Seizures
Entry into interphase
Alignment of chromosomes
Development of cerebral cortex
Differentiation of neurons
Development of neurons
Differentiation of nervous system
Organization of cytoplasm
Microtubule dynamics
Organization of cytoskeleton
Cell viability
Cell survival
Segregation of chromosomes
Mitosis
Cell cycle progression
Organismal death
-log10(FDR p-value)
Butterfly
Anxiety
Cell viability of neurons
Memory
Outgrowth of axons
Senescence of fibroblasts
Long-term potentiation
Potentiation of synapse
Senescence of connective tissue cells
Senescence of cells
Bundling of microtubules
Coordination
Cognition
Outgrowth of neurites
Outgrowth of neurons
Learning
Growth of axons
Alignment of chromosomes
Entry into interphase
Outgrowth of cells
Motor dysfunction or movement disorder
Movement Disorders
Formation of cellular protrusions
Growth of neurites
Quantity of neurons
Migration of neurons
Proliferation of neuronal cells
Neuritogenesis
Morphogenesis of neurons
Seizure disorder
Seizures
Segregation of chromosomes
Development of neurons
Organization of cytoplasm
Microtubule dynamics
Organization of cytoskeleton
-log10(FDR p-value)
z-score
-4 0 4
C
Senescence of connective tissue cells
Proliferation of connective tissue cells
Growth of connective tissue
Limb defect
Apoptosis of neurons
Outgrowth of axons
Death of embryo
Long-term potentiation
Bundling of microtubules
Colony formation
Growth of axons
Initiation of replication of DNA
Senescence of cells
Invasion of cells
Quantity of neurons
Neonatal death
Outgrowth of neurites
Cognition
Entry into interphase
Outgrowth of neurons
Learning
Motor dysfunction or movement disorder
Movement Disorders
Size of body
Alignment of chromosomes
Outgrowth of cells
Formation of cellular protrusions
Growth of neurites
Proliferation of neuronal cells
Migration of neurons
Cell viability
Seizure disorder
Cell survival
Seizures
Development of neurons
Organization of cytoplasm
Microtubule dynamics
Segregation of chromosomes
Organization of cytoskeleton
-log10(FDR p-value)
Peanut
Initiation of replication of DNA
Death of embryo
Tonic-clonic seizure
Generalized seizures
Convulsion
Outgrowth of axons
Apoptosis of neurons
Long-term potentiation
Potentiation of synapse
Senescence of cells
Growth of axons
Entry into interphase
Invasion of cells
Guidance of axons
Cognition
Quantity of neurons
Learning
Formation of cellular protrusions
Alignment of chromosomes
Cell movement of neurons
Migration of neurons
Neonatal death
Movement Disorders
Motor dysfunction or movement disorder
Growth of neurites
Proliferation of neuronal cells
Neuritogenesis
Seizure disorder
Perinatal death
Seizures
Development of neurons
Segregation of chromosomes
Organization of cytoplasm
Microtubule dynamics
Organization of cytoskeleton
-log10(FDR p-value)

**Figure S2.** ***Related to Figure 2.*** **Time-course heatmaps, PCA plots and the extended list of biological functions during hCO differentiation in all microwell and 96-well conditions.**

(A-C) n=3 biological replicates for each condition with 4-10 hCOs in each replicate.

A) Time-course heatmaps for all microwell shapes and 96-wells. Generated with CLC genomics workbench (version 20.0.2) using trimmed mean of M values (TMM) normalization method and Euclidean distance with complete linkage displaying a fixed number of 10,000 features with the highest coefficient of variance.

B) Time-course PCA of gene expression during hCO differentiation for all microwell shapes and 96-well conditions.

C) Extended list of biological functions between days 40 and 10 of hCO differentiation with z-score $\geq |2|$ and FDR p-value $< 0.01$ identified through IPA analysis. Differentially expressed genes with FDR p-value $< 0.05$, $\log_2$ (FC) $\geq |1.5|$ and max mean RPKM $\geq 10$ were used in this analysis. 809 (96-well), 877 (Round), 621 (Butterfly), 807 (Peanut) and 788 (C) genes met this criteria.

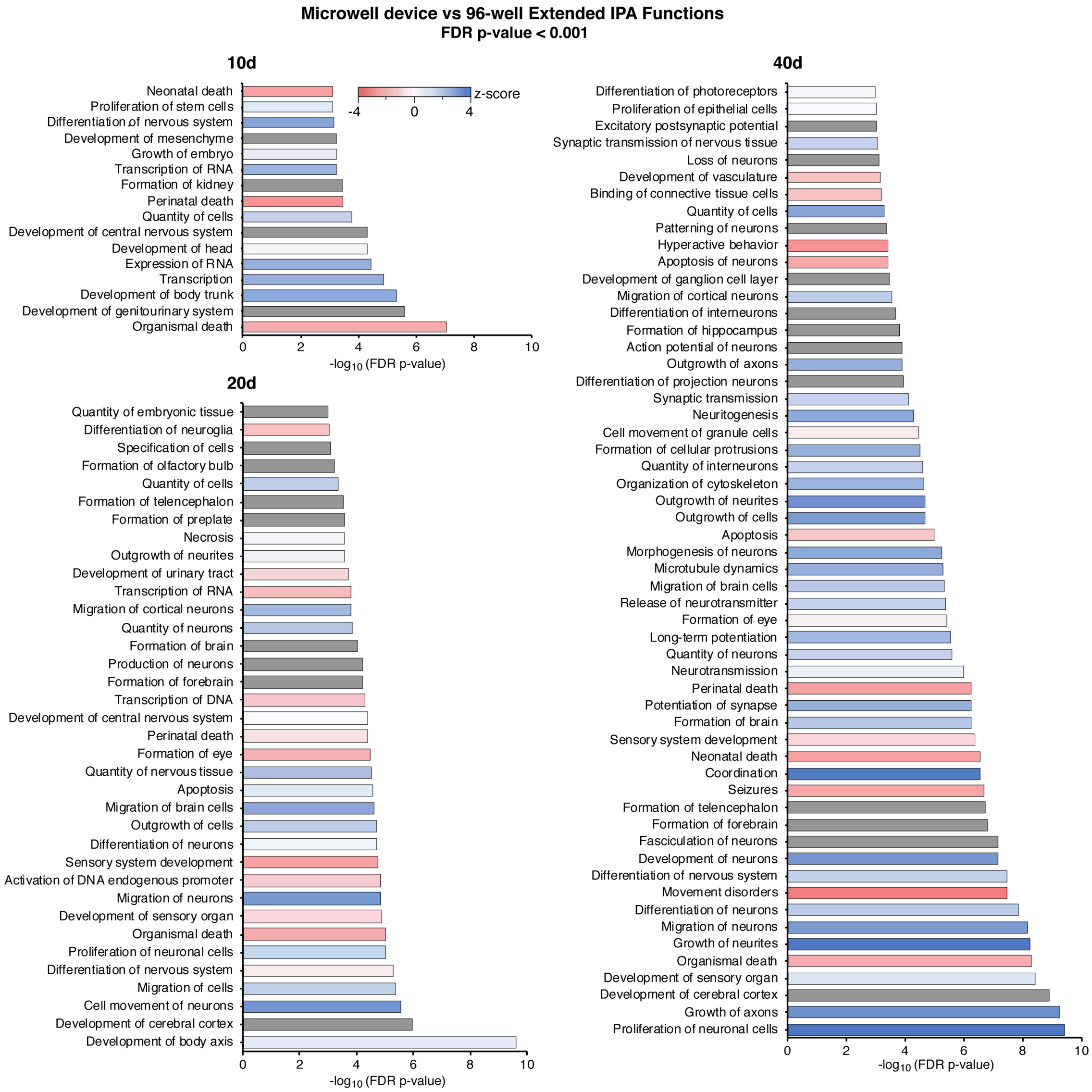


**Figure S3.** ***Related to Figure 3.*** **Extended list of biological functions between microwell device and 96-well plate.**

Functions with FDR p-value < 0.001 identified through IPA analysis are listed. Differentially expressed genes with FDR p-value < 0.05, $\log_2$ (FC) ≥ |1.5| and max mean RPKM ≥ 5 were used in this analysis. Gray bars indicate no z-score was calculated. n=12 biological replicates for microwell device (all shapes) with 4-8 hCOs in each replicate, n=3 for 96-well plate with 4-8 hCOs in each. 51 (day 10), 56 (day 20) and 166 (day 40) genes met this criteria.

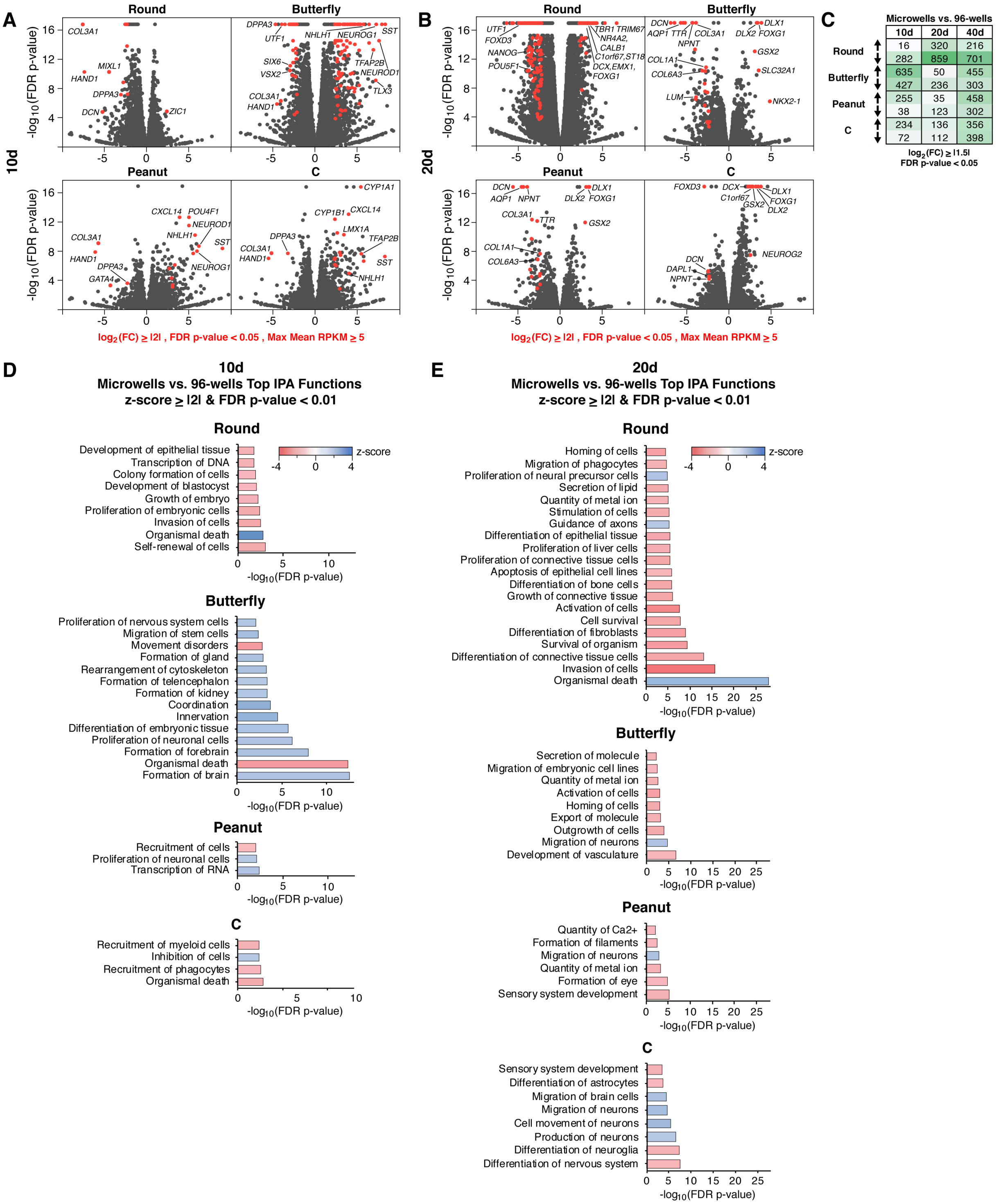

A
B
C
D
E
10d
20d
Round
Butterfly
Peanut
C
-log10(FDR p-value)
COL3A1
MIXL1
HAND1
DPPA3
DCN
ZIC1
DPPA3
UTF1
NHLH1
NEUROG1
SST
SIX6
VSX2
TFAP2B
NEUROD1
TLX3
CXCL14
POU4F1
GATA4
CYP1A1
CYP1B1
LMX1A
UTF1
FOXD3
NANOG
POU5F1
TBR1 TRIM67
NR4A2,
CALB1
C1orf67,ST18
DCX,EMX1,
FOXG1
AQP1
TTR
NPNT
COL1A1
COL6A3
LUM
DLX1
DLX2
GSX2
SLC32A1
NKX2-1
DAPL1
NEUROG2
DCX
C1orf67
log2(FC) ≥ |2| , FDR p-value < 0.05 , Max Mean RPKM ≥ 5
Microwells vs. 96-wells
10d 20d 40d
Round ↑ 16 320 216
Round ↓ 282 859 701
Butterfly ↑ 635 50 455
Butterfly ↓ 427 236 303
Peanut ↑ 255 35 458
Peanut ↓ 38 123 302
C ↑ 234 136 356
C ↓ 72 112 398
log2(FC) ≥ |1.5|
FDR p-value < 0.05
10d
Microwells vs. 96-wells Top IPA Functions
z-score ≥ |2| & FDR p-value < 0.01
Round
Development of epithelial tissue
Transcription of DNA
Colony formation of cells
Development of blastocyst
Growth of embryo
Proliferation of embryonic cells
Invasion of cells
Organismal death
Self-renewal of cells
z-score
Butterfly
Proliferation of nervous system cells
Migration of stem cells
Movement disorders
Formation of gland
Rearrangement of cytoskeleton
Formation of telencephalon
Formation of kidney
Coordination
Innervation
Differentiation of embryonic tissue
Proliferation of neuronal cells
Formation of forebrain
Organismal death
Formation of brain
Peanut
Recruitment of cells
Proliferation of neuronal cells
Transcription of RNA
C
Recruitment of myeloid cells
Inhibition of cells
Recruitment of phagocytes
Organismal death
20d
Microwells vs. 96-wells Top IPA Functions
z-score ≥ |2| & FDR p-value < 0.01
Round
Homing of cells
Migration of phagocytes
Proliferation of neural precursor cells
Secretion of lipid
Quantity of metal ion
Stimulation of cells
Guidance of axons
Differentiation of epithelial tissue
Proliferation of liver cells
Proliferation of connective tissue cells
Apoptosis of epithelial cell lines
Differentiation of bone cells
Growth of connective tissue
Activation of cells
Cell survival
Differentiation of fibroblasts
Survival of organism
Differentiation of connective tissue cells
Invasion of cells
Organismal death
Butterfly
Secretion of molecule
Migration of embryonic cell lines
Quantity of metal ion
Activation of cells
Homing of cells
Export of molecule
Outgrowth of cells
Migration of neurons
Development of vasculature
Peanut
Quantity of Ca2+
Formation of filaments
Migration of neurons
Quantity of metal ion
Formation of eye
Sensory system development
C
Sensory system development
Differentiation of astrocytes
Migration of brain cells
Migration of neurons
Cell movement of neurons
Production of neurons
Differentiation of neuroglia
Differentiation of nervous system

**Figure S4.** ***Related to Figure 4.*** **Volcano plots and top biological functions between individual microwell shapes and 96-wells at day 10 and 20.**

A) Volcano plots of differentially expressed genes between individual microwell shapes and 96-wells at 10 days. Red highlighted genes: FDR p-value < 0.05, $log_2$ (FC) ≥ |2| and max mean RPKM ≥ 5.

B) Volcano plots of differentially expressed genes between individual microwell shapes and 96-wells at 20 days. Red highlighted genes: FDR p-value < 0.05, $log_2$ (FC) ≥ |2| and max mean RPKM ≥ 5.

C) The number of differentially expressed genes between individual microwell shapes and 96-wells with FDR p-value <0.05 and $log_2$ (FC) ≥ |1.5|.

D) Top biological functions between individual microwell shapes and 96-wells at 10 days with z-score ≥ |2| and FDR p-value < 0.01 identified through IPA analysis. Differentially expressed genes with FDR p-value < 0.05, $log_2$ (FC) ≥ |1.5| and max mean RPKM ≥ 5 were used in this analysis. 49 (Round), 209 (Butterfly), 23 (Peanut) and 30 (C) genes met this criteria.

E) Top biological functions between individual microwell shapes and 96-wells at 20 days with z-score ≥ |2| and FDR p-value < 0.0 identified through IPA analysis. Differentially expressed genes with FDR p-value < 0.05, $log_2$ (FC) ≥ |1.5| and max mean RPKM ≥ 5 were used in this analysis. 49 (Round), 209 (Butterfly), 23 (Peanut) and 30 (C) genes met this criteria. 453 (Round), 51 (Butterfly), 35 (Peanut) and 38 (C) genes met this criteria.

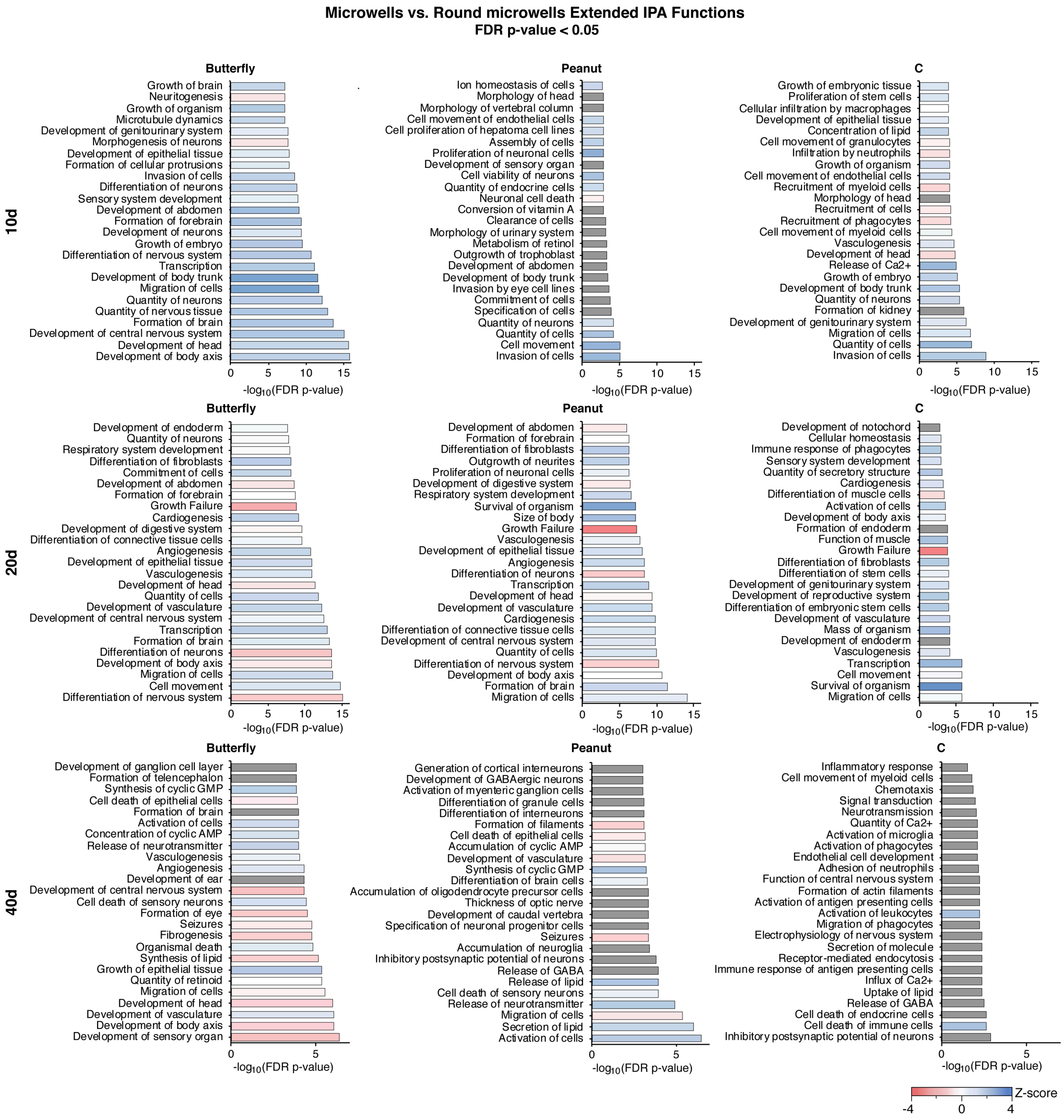


**Figure S5.** ***Related to Figure 5.*** **Extended list of biological functions between individual microwell shapes and round microwells.**

Functions with FDR p-value < 0.05 identified through IPA analysis. Differentially expressed genes with FDR p-value < 0.05, $\log_2$ (FC) ≥ |1.5| and max mean RPKM ≥ 5 were used in this analysis. 149 (Butterfly), 28 (Peanut) and 35 (C) genes met this criteria in day 10 hCOs. 286 (Butterfly), 266 (Peanut) and 88 (C) genes met this criteria in day 20 hCOs. 106 (Butterfly), 69 (Peanut) and 10 (C) genes met this criteria in day 40 hCOs. Gray bars indicate no z-score was calculated. n=3 biological replicates for each condition with 4-10 hCOs in each replicate.
